# Shear-Mediated Platelet Microparticles Demonstrate Phenotypic Heterogeneity as to Morphology, Receptor Distribution, and Hemostatic Function

**DOI:** 10.1101/2023.02.08.527675

**Authors:** Yana Roka-Moiia, Kaitlyn Ammann, Samuel Miller-Gutierrez, Jawaad Sheriff, Danny Bluestein, Joseph E. Italiano, Robert C. Flaumenhaft, Marvin J. Slepian

## Abstract

**Objective:** Implantable cardiovascular therapeutic devices (CTD) including stents, percutaneous heart valves and ventricular assist devices, while lifesaving, impart supraphysiologic shear stress to platelets resulting in thrombotic and bleeding device-related coagulopathy. We previously demonstrated that shear-mediated platelet dysfunction is associated with downregulation of platelet GPIb-IX-V and αIIbβ3 receptors via generation of platelet-derived microparticles (PDMPs). Here, we test the hypothesis that shear-generated PDMPs manifest phenotypical heterogeneity of their morphology and surface expression of platelet receptors, and modulate platelet hemostatic function.

**Approach and Results:** Human gel-filtered platelets were exposed to continuous shear stress and sonication. Alterations of platelet morphology were visualized using transmission electron microscopy. Surface expression of platelet receptors and PDMP generation were quantified by flow cytometry. Thrombin generation was quantified spectrophotometrically, and platelet aggregation in plasma was measured by optical aggregometry. We demonstrate that platelet exposure to shear stress promotes notable alterations in platelet morphology and ejection of several distinctive types of PDMPs. Shear-mediated microvesiculation is associated with the differential remodeling of platelet receptors with PDMPs expressing significantly higher levels of both adhesion (α_IIb_β_3_, GPIX, PECAM-1, P-selectin, and PSGL-1) and agonist-evoked receptors (P_2_Y_12_ & PAR1). Shear-mediated PDMPs have a bidirectional effect on platelet hemostatic function, promoting thrombin generation and inhibiting platelet aggregation induced by collagen and ADP.

**Conclusions:** Shear-generated PDMPs demonstrate phenotypic heterogeneity as to morphologic features and defined patterns of surface receptor alteration, and impose a bidirectional effect on platelet hemostatic function. PDMP heterogeneity suggests that a range of mechanisms are operative in the microvesiculation process, contributing to CTD coagulopathy and posing opportunities for therapeutic manipulation.

## INTRODUCTION

Implantable cardiovascular therapeutic devices (CTD) have become the mainstay of advanced therapy for a broad range of cardiovascular diseases. In recent years, we have witnessed growth in implantation of stents for obstructive atherosclerotic disease in a myriad of anatomic locations,^1–4^ percutaneous heart valves and restorative devices (e.g. TAVR and mitral clips),^5–8^ novel sensing systems, and increased utilization of mechanical circulatory support (MCS), with placement of percutaneous or surgical ventricular assist devices for acute and chronic advanced and end-stage heart failure.^9–12^ While all of these devices have demonstrated hemodynamic and functional efficacy, implantation still comes at a cost in terms of hemocompatibility.^13–15^ In an attempt to enhance device hemocompatibility, anticoagulant and antiplatelet agents are administered clinically.^16–18^ While pharmacologic agents have demonstrated efficacy in limiting stent thrombosis,^18, 19^ for other devices, it is increasingly evident that drugs have limited efficacy and often contribute to increased bleeding adverse events.^20–23^ For both coronary stents and MCS, presently there is a move afoot to curtail or eliminate the use of antithrombotic agents to reduce bleeding.^16, 18, 20, 24, 25^

Hemocompatibility of implanted devices is mediated by a delicate balance of materials and surface considerations,^26, 27^ thrombogenicity and inflammatory potential of blood,^28, 29^ and flow-related disturbances imparted by abnormal shear stresses and turbulence.^30–32^ In recent years, device hemocompatibility has improved somewhat, though due to design-dictated flow regimes and geometries, supraphysiologic shear stress is still imparted to blood cells and proteins as they traverse devices with resultant biological consequences.^31, 33, 34^ We and others have reported on the role of elevated shear stress as a driver of an underlying device-related coagulopathy with both prothrombotic and bleeding features.^35–37^ Shear-mediated platelet activation via mechanotransductive and mechanodestructive means has been demonstrated to drive thrombosis.^35, 36, 38–40^ Concomitantly, continuous platelet exposure to hypershear results in acquired platelet dysfunction which contributes to device-related bleeding.^36, 37, 41–43^

Platelets are small anucleate blood cells essential for maintaining primary and secondary hemostasis.^44, 45^ Platelets operate in delicate equipoise between thrombosis and bleeding, biochemically and physiologically controlled to avoid inadvertent thrombosis or bleeding with dire systemic consequences. When exposed to biochemical and mechanical stimuli, platelets undergo activation, rapid shape change, release their secretory granules, and ultimately aggregate to form primary thrombi.^46, 47^ In their ability to sense the extracellular environment and interact with other platelets and cells, platelets rely on two classes of glycoproteins (GP) receptors: *adhesion* receptors and *agonist-evoked* receptors. Platelet *adhesion receptors* are abundantly expressed on the membrane surface, either constitutively (GPIb-V-IX, integrin α_IIb_β_3_, GPVI, PECAM-1)^46, 48, 49^ or “on-demand” as a result of platelet activation (P-selectin, PSGL-1).^50, 51^ During platelet aggregation, GPIb-V-IX primarily binds to von Willebrand factor (vWF), integrin α_IIb_β_3_ binds to fibrinogen and vWF as its major ligands, and GPVI interacts with collagen.^52–54^ PECAM-1, P-selectin, and PSGL-1 are responsible for platelet binding to leukocytes and vascular endothelium.^48, 50, 55^ The most well-known and therapeutically relevant *agonist receptors* are ADP-evoked P_2_Y_12_ and thrombin-evoked PAR1 receptors. These receptors are scarcely, yet constitutively, expressed on the platelet surface, and due to their potency in driving platelet activation, are targets for common antiplatelet agents.^56–58^ The surface expression of platelet GP receptors is tightly regulated to limit spontaneous platelet adhesion and activation upon interaction with plasma proteins and other cells. Enzymatic shedding of a ligand-binding ectodomain is the most well-documented regulatory mechanism of platelet receptor surface expression.^45, 59, 60^ Both continuous and activation-induced types of enzymatic shedding have been described for GPIb, GPVI, P-selectin, and PECAM-1.^45, 61, 62^ An alternative mechanism of downregulation of receptor surface expression is the restructuring of the entire platelet membrane as a result of microvesiculation and generation of microparticles.^37, 63^

Platelet-derived microparticles (PDMPs) are submicron, membrane-enclosed, extracellular vesicles released from platelets during their activation by strong agonists, aging, and apoptosis.^64–66^ Platelets and megakaryocytes are considered to be the primary sources of microvesicles circulating in blood, with up to 90% carrying canonical platelet antigens – CD41, CD42b, GPVI, and externalized phosphatidylserine.^67, 68^ Detailed mechanisms of PDMP formation have been described emphasizing the importance of high intracellular calcium, actin cytoskeleton rearrangement, and apoptotic machinery in this process.^69, 70^ Studies of PDMP proteomics have revealed several types of microvesicles differing in size, protein composition, and their effect on clotting and endothelial cell phenotype.^71–73^ PDMPs not only provide an additional surface for activation of the coagulation cascade and thrombin generation, but also deliver platelet receptors, nucleic acids, signaling and pro-inflammatory molecules to other cells, serving as a specialized mechanism of intravascular and extravascular cell-cell communication.^64, 74, 75^ Our research group and others have demonstrated that extended exposure to MCS-related shear stress promotes downregulation of platelet adhesion receptors and is associated with extensive microparticle generation *in vitro,* and *in vivo*.^37, 76–79^ Elevated levels of circulating microparticles were more pronounced in device-implanted patients experiencing post-implantation bleeding events as opposed to non-bleeders.^43, 79, 80^ For other CTDs, elevated levels of microparticles were also associated with hemostatic dysfunction, inflammation, and adverse events.^81, 82^ Despite the significance of PDMPs in the pathophysiology and diagnosis of device-related coagulopathy, their detailed phenotypic characterization and modulatory effect on platelet hemostatic function have not been investigated.

Here, we hypothesized that (1) exposure to supraphysiologic shear stress promotes alterations of platelet morphology, accompanied by generation of microparticles; (2) shear-mediated PDMPs manifest phenotypical heterogeneity of their morphology and surface expression of platelet *adhesion* receptors and *agonist-evoked* receptors; and (3) shear-mediated PDMPs modulate platelet hemostatic function. Specifically, we examined the effect of shear stress accumulation on morphology of platelets and shear-mediated PDMPs using transmission electron microscopy (TEM). We defined the effect of shear stress on the distribution of adhesion receptors and agonist-evoked receptors on platelets and microparticles using multi-colored flow cytometry. Finally, we tested the effect of shear-mediated PDMPs on platelet hemostatic function, i.e. thrombin generation and agonist-induced platelet aggregation.

## METHODS

### Blood collection and platelet isolation

Blood was obtained via venipuncture from healthy, volunteers following their written consent. The study protocol was approved by the IRB of the University of Arizona (protocol #1810013264). Freshly collected blood was anticoagulated with acid citrate dextrose solution at a ratio of 9:1. Platelet-rich plasma (PRP) was obtained by centrifugation of anticoagulated blood at 400*g* for 15 min. Residual blood was centrifuged at 1430*g* for 20 min, yielding platelet-poor plasma. Gel-filtered platelets (GFP) were isolated from PRP by gel chromatography through Sepharose-2B.^83^ Platelet fractions were stored and handled at room temperature to minimize storage-associated platelet function decline.^84^ Similarly, to limit 37°C-stimulated platelet apoptosis^85, 86^ and associated alterations of the surface expression of platelet receptors^45^ and microparticle generation,^87^ most experiments were performed at room temperature if not otherwise specified.

### Platelet exposure to shear stress and sonication

Recalcified GFP (20,000 platelets/μL, 2.5 mM CaCl_2_) was subjected to shear stress in a hemodynamic shearing device, a computer-controlled modified cone-plate-Couette viscometer allowing application of uniform continuous shear stress of defined magnitude (10, 30, 50, and 70 dyne/cm^2^).^88^ For these conditions, the shear stress accumulation over time represents repeated platelet passages through high-shear regions of MCS devices, and was selected based on our previous numerical studies of the hemodynamics of MCS devices as compared with physiological levels of shear stress present within the normal circulation.^89, 90^ For shear exposure experiments, 10 min was standardized as shear exposure time based on our previous reports.^36, 37, 89, 91^ Alternatively, recalcified GFP (20,000 platelets/μL, 2.5 mM CaCl_2_) were subjected to sonication for 10s at 50% of power (Branson Ultrasonics™ SLPt Sonifier).

To generate PDMPs by shear stress, recalcified GFP (100,000 platelets/μL, 2.5 mM CaCl_2_) were subjected to 70 dyne/cm^2^ shear stress for 30 min. Alternatively, recalcified GFP (100,000 platelets/μL, 2.5 mM CaCl_2_) were subjected to sonication as described above. The sheared and sonicated samples were centrifuged twice at 2000g for 10 min to sediment platelets.^92^ Then, the supernatant containing PDMPs was collected and stored on ice until used as a PDMP-rich fraction.

### Platelet and microparticle imaging via transmission electron microscopy

Following shear exposure or sonication, GFP (100,000 platelets/μL, 2.5 mM CaCl_2_) were fixed with 2.5% (v/v) formaldehyde, 2.5% (v/v) glutaraldehyde in 0.1 M sodium cacodylate buffer, pH 7.4, and incubated at room temperature for 30 min. Fixed GFP were then centrifuged at 1500g for 15 min to sediment intact platelets (platelet-rich low-speed fraction). The supernatant was collected and centrifuged at 20,000g for 30 min to obtain a PDMP-rich pellet (high-speed fraction). Platelet and PDMP-rich pellets were stored separately in 0.2 M sodium cacodylate buffer at 4°C until TEM imaging.^93^

Prior to TEM imaging, platelet- and PDMP-rich pellets were washed in 0.1 M sodium cacodylate buffer and fixed with 1% osmium tetroxide (OsO_4_)/1.5% potassium ferrocyanide (KFeCN_6_) for 1 hour at room temperature. Fixed pellets were washed in water, 0.2 M maleate buffer, pH 5.15, and stained with 1% (v/v) uranyl acetate in maleate buffer for 1 hour. Samples were then dehydrated in increasing grades of ethanol (10 minutes each: 50%, 70%, 90%, 100%) and infiltrated on a 1:1 mixture of propylene oxide and TAAB resin overnight at 4°C. Samples were then embedded in TAAB Epon and polymerized at 60°C for 48 hours. Ultrathin sections (60 nm) of pellets were cut utilizing a Reichert Ultracut-S microtome (Leica Microsystems, Bannockburn, IL), transferred onto copper grids, and stained with lead citrate. Sections were examined in a JEOL 1200 EX (Tokyo, Japan) or TecnaiG^2^ Spirit BioTWIN transmission electron microscopes (Hillsboro, OR) at an accelerating voltage of 80 kV. Images were recorded with an AMT 2k CCD camera.

### Surface expression of platelet receptors on platelets and microparticles

Flow cytometric detection of platelet surface receptors was performed following published recommendations.^94, 95^ Briefly, 100 μL of sheared or sonicated GFP (20,000 platelets/μL, 2.5 mM CaCl_2_) were double-stained with five fluorescein-conjugated antibody pairs as follows: anti-CD41-APC (clone MEM-6,1:10) & anti-PAR1-AF488 (clone 731115, 1:20), anti-CD42a-FITC (clone GR-P, 1:250) & anti-CD62P-APC (clone Psel.KO2.3, 1:80), anti-CD31-FITC (clone WM-59, 1:250) & anti-PSGL1-PE (clone KPL-1, 1:5), anti-CD41-APC (clone MEM-6,1:10) & annexin V-FITC (1:20), anti-GP6-PE (clone HY101, 1:250) & anti-P2RY12-FITC (clone S16001E, 1:20). Samples were incubated for 30 min and then fixed with 2% PFA (filtered via 0.22 µm syringe filter) for 20 min. Stained and fixed platelets were diluted up to 1 mL with 50 mM PBS (pH 7.4, filtered via 0.22 µm syringe filter). Flow cytometry was conducted on a FACSCanto II flow cytometer (BD Biosciences, San Jose, CA). Ten thousand events were captured within the stopping gate “Platelet + Microparticles”. FCS Express Version 3 software (DeNovo Software, San Jose, CA) was applied to analyze flow cytometry data. Single platelets were distinguished from microparticles based on their forward versus side scatter characteristics (FSC-A/SSC-A), as compared with standard polystyrene beads of 880 nm and 1350 nm size from the SPHERO^TM^ Nano Fluorescent particle standard kit (Spherotech, Lake Forest, IL).^37^ Marker-positive platelet and microparticle count were expressed as % of all events captured in a joint gate “Platelets + Microparticles”. Platelet and microparticle size was assessed as their median forward scatter (MFS).^96, 97^ The arbitrary density of surface receptors on platelet and microparticle surfaces was calculated as the median fluorescence intensity (MFI) of the population normalized to the median FSC indicating platelet or microparticle size.^37, 98^

### Thrombin generation on platelets and microparticles

The effect of platelets and PDMPs on prothrombin activation by factor Xa was assessed using the modified thrombin generation assay.^99^ Following platelet exposure to shear stress, sonication, or PDMP-rich fraction, 100 μL of recalcified GFP (5,000 platelets/μL, 5 mM CaCl_2_) were incubated with 200 nM acetylated prothrombin and 100 pM factor Xa (Enzyme Research Laboratories, South Bend, IN) in 20 mM HEPES buffer (pH 7.4), containing 130mM NaCl and 0.1% BSA at 37°C for 10 min. Then, 10 μL of each GFP sample were tested for thrombin activity in microplate wells, containing 0.3 mM Chromozym TH (Roche Diagnostics GmbH), 3mM EDTA. Kinetic changes in light absorbance (A405) were recorded for 7 min using a Versa MAX microplate reader (Molecular Devices Corp., Sunnyvale, CA). The rate of thrombin generation was calculated as a slope of the kinetic curve (1/min) using the SoftMax Pro6 software (Molecular Devices Corp).

### Platelet aggregation induced by biochemical agonists

Platelet aggregation was measured following PRP exposure to PDMPs and biochemical agonists. Prior to every aggregation test, PRP was diluted to 100,000 platelets/μL with modified Tyrode’s buffer and recalcified with 1 mM CaCl_2_. Then, 300 µL of PRP was transferred into an aggregation cuvette and incubated with 30 μL of PDMP-rich fraction (or Tyrode’s buffer for vehicle control) for 10 min at 37°C with stirring. Platelet aggregation was initiated by the following biochemical agonists: 2 ug/mL collagen, 10 μM ADP, or 5 μM TRAP-6. Alternatively, platelet aggregation was induced by the PDMP-rich fractions alone, and no biochemical agonist was added. The relative changes in light transmittance were recorded for 10 min by the optical aggregometer PAP-8E (BioData Corporation, Horsham, PA). The area under the curve was calculated by the PAP-8 software (BioData Corporation, Horsham, PA).

### Statistical analysis

Results from 4 to 7 independent experiments with different donors were summarized in figures. All flow cytometry and aggregometry samples were run in duplicate. The data were tested for normality using the Shapiro-Wilk normality test, and then statistically analyzed using one-way analysis of variance (ANOVA) followed by Dunnett’s multiple comparisons test and paired t-test for paired comparisons all from GraphPad Prism 8 software (GraphPad Software Inc., San Diego, CA). Averages are reported as the Mean ± SEM. The level of statistical significance is indicated in the figures as *p* ˂ 0.05 and *p* ˂ 0.01.

## RESULTS

Herein, we performed a complex phenomenological study of platelets and PDMPs generated under mechanical shear stress conditions. First, using TEM, we visualized alterations of platelet morphology accompanying PDMP generation during shear stress and sonication. Second, using immunostaining and quantitative multi-colored flow cytometry, we characterized the surface distribution of eight surface receptors on platelets and microparticles: 1) constitutively expressed adhesion receptors (αIIbβ3, GPIX, GPVI, PECAM-1), 2) inducible adhesion receptors (P-selectin, PSGL1), and 3) agonist-evoked receptors (P2Y12, PAR1). Lastly, we analyzed the effect of PDMPs generated by shear stress and sonication on platelet hemostatic function, namely aggregation and thrombin generation.

### 1.1. Shear-induced alterations of platelet morphology and microparticle generation

Transmission electron microscopy was utilized to reveal morphologic details of platelets and microparticles following exposure to mechanical stress via shear stress or sonication.^93^ Following stress exposure, platelets were sequentially centrifuged at a low speed (1500*g*) to pellet platelets, and at high-speed (20,000*g*) to pellet microparticles and any residual platelets. Platelet- and microparticle-rich fractions were then examined via TEM.

The TEM examination of the *low-speed fraction* revealed morphological details of resting, sheared, and sonicated platelets (**Fig. 1A-D**). Resting platelets were generally round, ovoid, or ellipsoidal with smooth contours and continuous membranes, containing uniform cytoplasm with multiple discrete granules (**Fig. 1A**). With increasing shear magnitude, platelet morphology remained largely intact through exposure to 30 dyne/cm^2^ (data not shown). At 50 dyne/cm^2^ shear, a moderate degree of platelet size alteration, pseudopod extension, and microparticle formation was detected (**Fig. 1B**); at 70 dyne/cm^2^ shear, significant alteration of platelet morphology was evident and widespread (**Fig. 1C**). More than half of platelets observed per field had significantly altered morphology – being either round with pale, depleted cytoplasm or fragmented with noticeable microvesiculation and emerging free PDMPs. Other platelets appeared smaller, with early signs of pseudopod formation. Sonication, as a form of mechanical shear generated via acoustic cavitation, largely disrupted platelets with residual ghost-like structures and free PDMPs (**Fig. 1D**).

**Figure 1.**
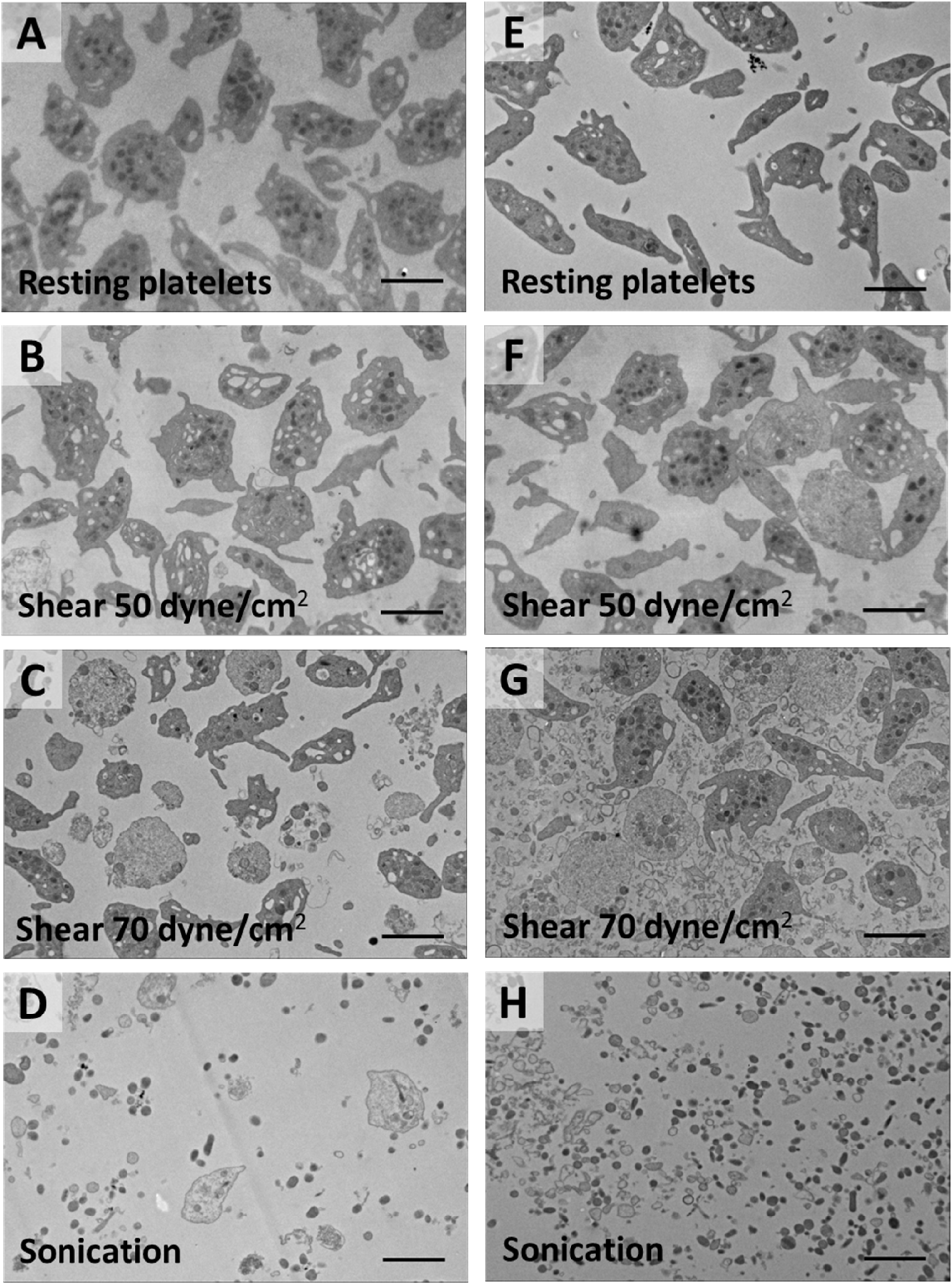
Shear stress of increasing magnitude and sonication promote alterations of platelet morphology & generation of platelet-derived microparticles. Illustrative transmission electron microscopy images of (A-D) platelets and (E-H) platelet-derived microparticles following exposure to shear stress (50 & 70 dynes/cm^2^) and sonication. (A-D) Platelets were pelleted via low-speed centrifugation (1,500*g*, 15 min). (E-H) Platelets and microparticles were pelleted by high-speed centrifugation (20,000*g*, 30 min). Scale bar represents 2 μm. Direct magnification: 1200× (A), 2900× (B, F), and 3000× (C-H).

The microparticle-rich fraction obtained by *high-speed* centrifugation of resting platelets contained some intact platelets and platelet fragments not pelleted by low-speed centrifugation along with traces of microparticles (**Fig. 1E-H**). Resting platelets had morphology similar to the low-speed fraction, though were of lower number (**Fig. 1E**). Rare cigar-shaped platelet fragments (100 x 800 nm) were visible, typically devoid of granules with cytoplasmic density similar to intact platelets. With an increase of shear magnitude to 50 dyne/cm^2^, a mild increase in the number of platelet fragments and PDMPs was detected (**Fig. 1F**). Notably, at 70 dyne/cm^2^ shear, a significant alteration in platelet morphology and a prominent increase of PDMPs was obvious (**Fig. 1G**). Intact platelets were outnumbered by residual platelets with loss of cytoplasmic density, being depleted of their granules. Following sonication, no intact platelets were observed, rather only microparticles (**Fig. 1H**). Sonicated PDMPs appeared as circular, ellipsoidal, or amorphous particles with a range of densities; some were similar to platelet granules and granules devoid of contents.

The range of platelet and PDMP shapes and densities after exposure to 70 dyne/cm^2^ shear stress is best seen in **Figure 2A**. Platelet membrane thinning and fragmentation were evident (**Fig. 2B-C**). Two types of PDMPs were noted: 1) large irregular particles, likely fragments of platelets or platelet structures evident in **Fig. 2A-G**; and 2) small circular or ellipsoidal particles – “granule-like”, with evident contrast density, or devoid of contents, best seen in **Fig. 2F-G**. Platelet granules were proximate to plasma membrane, with apparent budding from the membrane (**Fig. 2E**).

**Figure 2.**
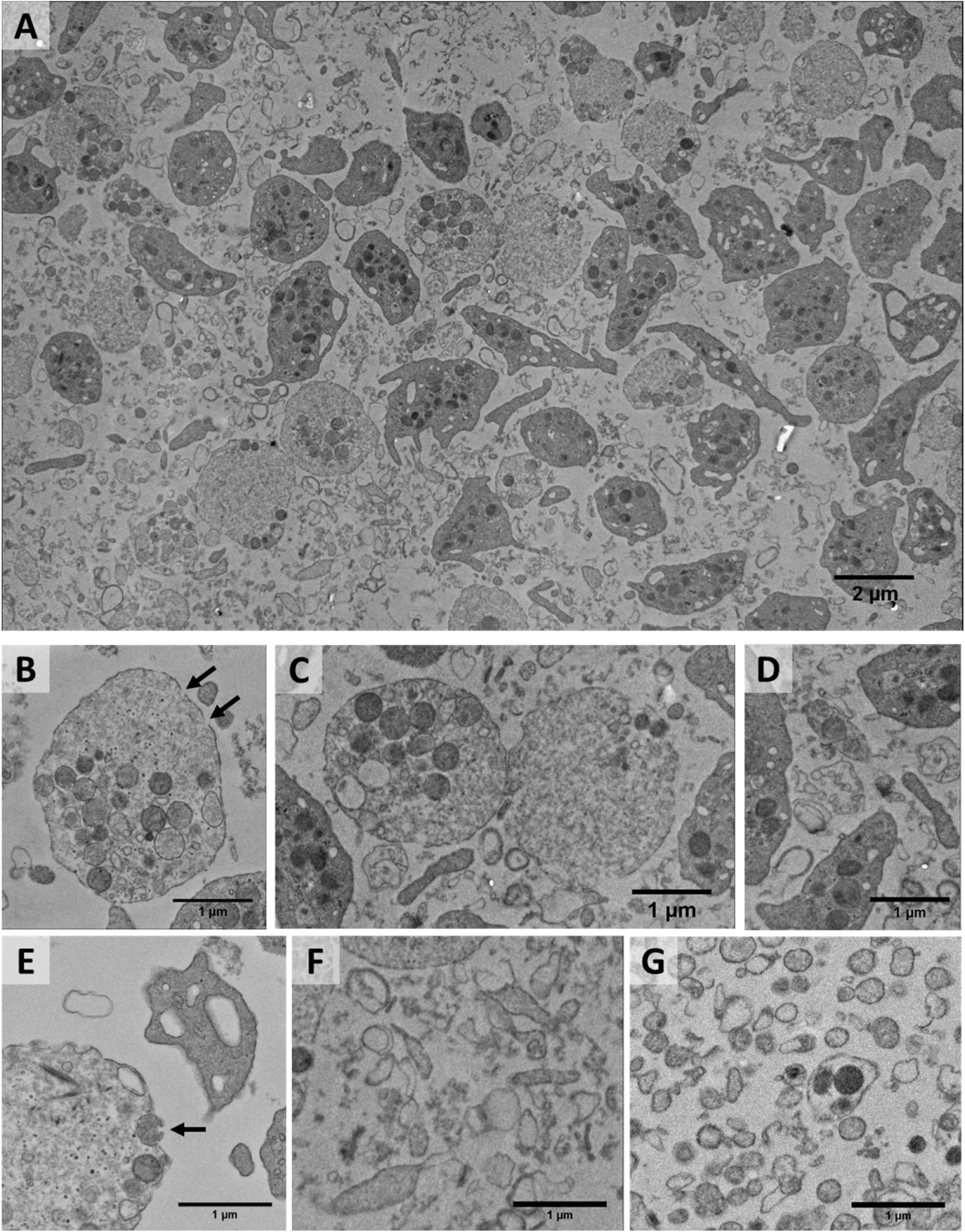
Illustrative thin-section electron microscopy images of platelets and microparticles following exposure to 70 dyne/cm^2^ shear stress. Microparticles were pelleted by high-speed centrifugation (20,000*g*, 30 min). Scale bar represents 1 μm. Direct magnification: 3000× (A-G).

The number of microparticles generated by shear stress and sonication was also assessed. Two broad groups of PDMPs were analyzed: 1) small particles ranging from 150 - 500 nm and 2) large particles ranging from 500 - 1000 nm. Small platelet fragments and debris were excluded from the analysis. An evident increase in the small particle population occurred following platelet exposure to high shear stress (**Fig. S1**). At 70 dyne/cm^2^ shear, 113+3 particles/3000× field were noted (ANOVA: *p*<0.01 vs. Unsheared), as compared with 26±4, 23±4, 37±10, and 22±1 particles/3000× field, for 50, 30, 10 dyne/cm^2^ and no shear, respectively. Sonication resulted in even greater PDMP generation: 551±381 particles/3000× field (ANOVA: *p*<0.01 vs. Unsheared). For larger particles, 37±4 particles/3000× field were observed at 70 dyne/cm^2^ shear, as compared with 7±1, and 3±1 particles/3000× field at 50 and 30 dyne/cm^2^ shear, respectively (data not shown). After 10 dyne/cm^2^ shear, no large particles were evident.

### 1.2. αIIbβ3 and GPIX distribution on platelets and platelet-derived microparticles

Platelet exposure to increasing shear stress and sonication resulted in a significant increase in PDMP and decrease in platelet populations. A set of typical dot diagrams captured by flow cytometry is shown in **Figure 3**. The CD41+ PDMP population significantly increased with the magnitude of shear (**Fig. 3B-E**, gate “Microparticles”); similarly, sonication resulted in dramatic microvesiculation and a prominent increase of PDMPs (**Fig. 3F**). As indicated in **Figure 4A-B**, a statistically significant increase in both CD41+ and CD42a+ PDMP populations was observed after 30 dyne/cm^2^ shear, and at 70 dyne/cm^2^ shear, CD41+ and CD42a+ PDMPs accounted for 9.1±1.2% and 5.7±0.4% of all events, respectively. Sonication was even more powerful, generating numerous CD41+ and CD42a+ PDMPs: 28.6±1.8% and 14.8±1.2%, respectively. At the same time, both CD41+ and CD42a+ *platelet* populations decreased gradually with an increase in shear stress magnitude (**Fig. 4C-D**). Sonication resulted in an even greater decrease of platelet count: 54.3±2.9% vs. 95.3±0.4% in the “No shear” group for CD41+ platelets, and 58.9±5.9% vs. 95.5±0.6% in the “No shear” group for CD42a+ platelets.

**Figure 3.**
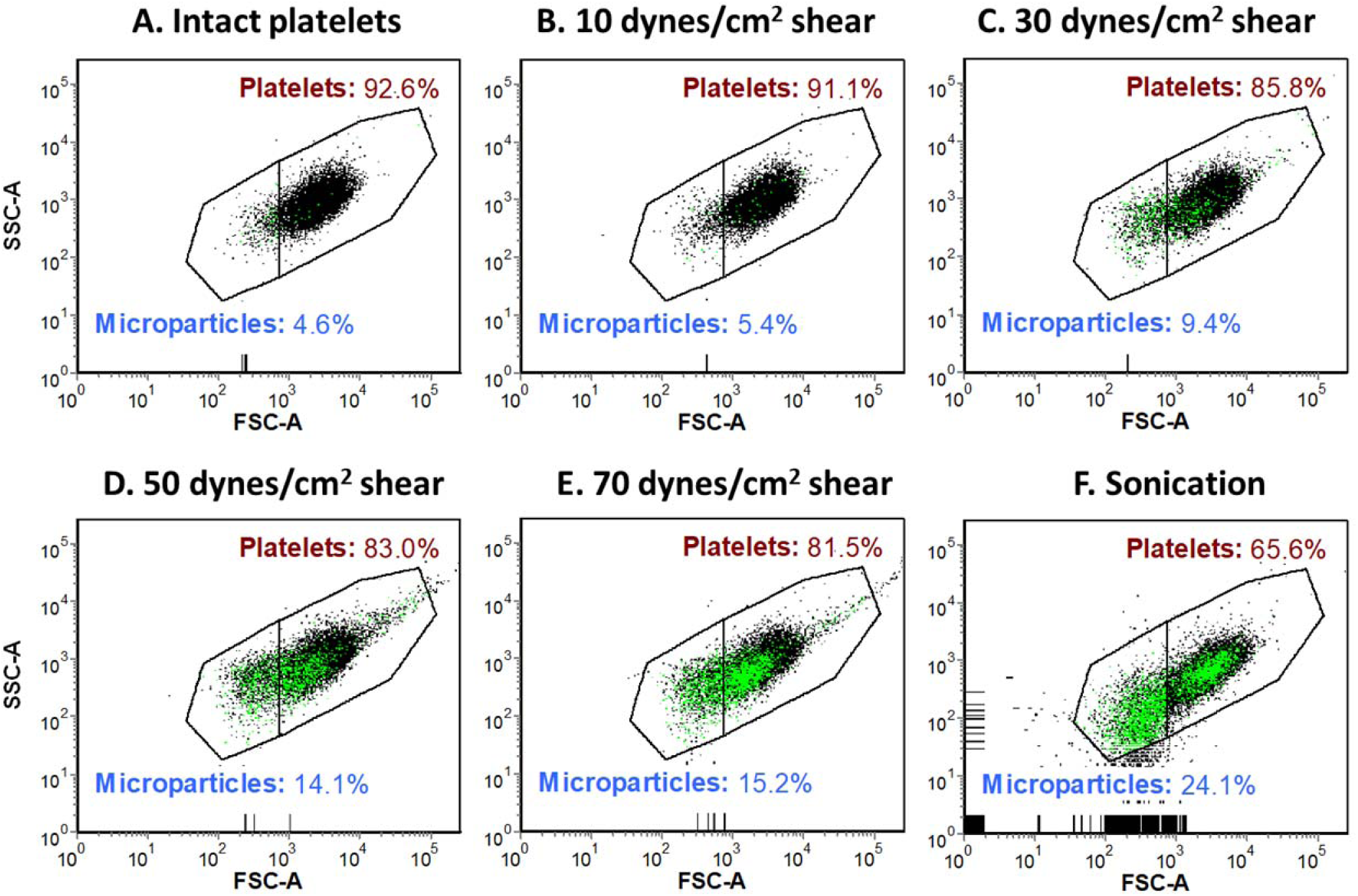
Platelet exposure to shear stress and sonication induces increased generation of platelet-derived microparticles (PDMPs). Illustrative dot diagrams of αIIb+ platelets and αIIb+ microparticles gated based on their forward and side scatter characteristics (A-F): A – intact platelets, B-E – sheared platelets, F – sonicated platelets. Black dots – CD41 positive platelets or microparticles, green dots – CD41+/annexin V+ double positive microparticles. FSC-A – forward scatter amplitude, SSC-A – side scatter amplitude.

**Figure 4.**
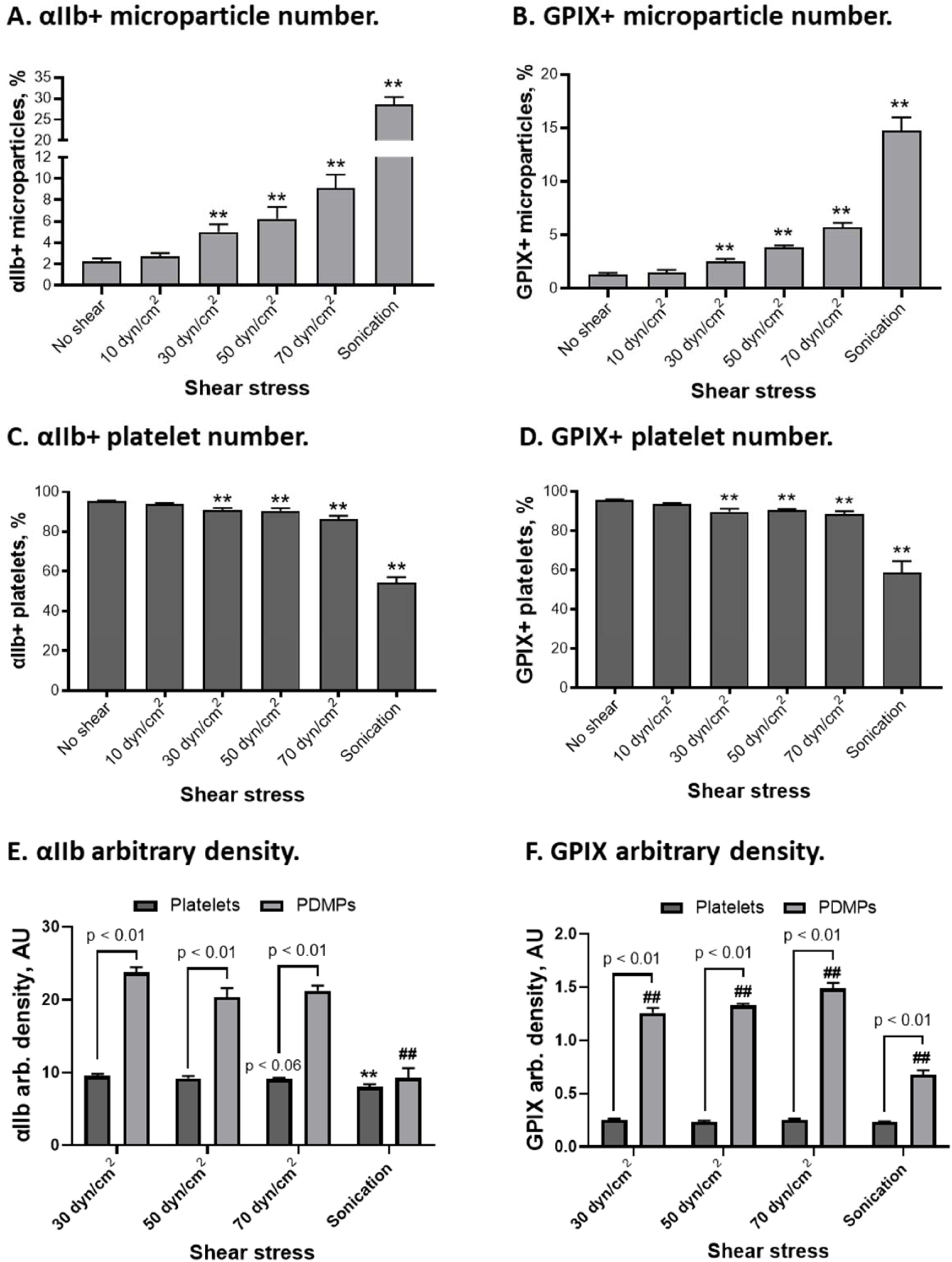
Distribution of αIIbβ3 integrin and GPIX on platelets and platelet-derived microparticles (PDMPs): A, B – αIIb (CD41)+ and GPIX (CD42a)+ PDMP number, C,D – αIIb (CD41)+ and GPIX (CD42a)+ platelet number, E, F – arbitrary density of αIIbβ3 (CD41) and GPIX (CD42) on platelets and PDMPs. N = 4-6. Mean ± SEM, 1-way ANOVA followed by Dunnett multiple comparisons test: **, ## - *p*<0.01 vs no shear for platelets and PDMPs.

Analyzing receptor distribution on platelet and microparticle surfaces, we (1) monitored alteration of the receptor density across all shear stress conditions and (2) compared the receptor density on platelets and microparticles. The arbitrary density of receptor distribution on the platelet surface was calculated by normalizing the fluorescence intensity (MFI) of platelet or microparticle to their forward scatter (MFS), a well-known flow cytometric parameter indicating particle size.^37, 98^ In **Figures S2 & S3**, alterations of fluorescence intensity, size, and receptor density with shear exposure or sonication are reported. Shear stress did not largely affect CD41 and CD42a fluorescence intensity (**Fig. S2A** & **S3A**) and platelet size (**Fig. S2B** & **S3B**). As such, the arbitrary density of α_IIb_β_3_ and GPIX receptors on platelets was not significantly altered (**Fig. S2E & S3E**, “Shear stress”). In contrast, sonication resulted in a decrease in platelet size and fluorescence intensity for both CD41 and CD42a, thus rendering a significant decrease in receptor arbitrary density, especially noticeable for α_IIb_β_3_. Similarly, the microparticle size and fluorescence intensity remained unchanged across all shear conditions for both receptors, while sonication generated slightly smaller PDMPs with nearly two-fold decreased surface receptor density as compared with shear-generated PDMPs (**Fig. S2D, S2F, S3D, S3F**, “Sonication”).

We also noticed that the arbitrary density of α_IIb_β_3_ and GPIX on *sheared* PDMPs was significantly higher than on platelets across all shear conditions (**Fig. 4E-F**, “Shear stress”). After 70 dyne/cm^2^ shear, CD41+ PDMPs expressed 2.3× higher receptor density than CD41+ platelets, while CD42a+ PDMPs showed 6× higher receptor density than CD42a+ platelets. In contrast, PDMPs generated by *sonication* demonstrated similar or barely elevated receptor density as compared to sonicated platelets (**Fig. 4E-F**, “Sonication”).

### 1.3. GPVI and PECAM-1 distribution on platelets and platelet-derived microparticles

The number of GPVI+ PDMPs was not elevated, even with the highest levels of shear applied (**Fig. 5A**), while the PECAM-1+ PDMP population increased gradually along with the shear magnitude and accounted for 6.2±0.6% after 70 dyne/cm^2^ shear (**Fig. 5B**). Sonication generated significantly greater amounts of GPVI+ and CD31+ microparticles, accounting for 6.8±1.2% and 18.9±0.6% of events, respectively.

**Figure 5.**
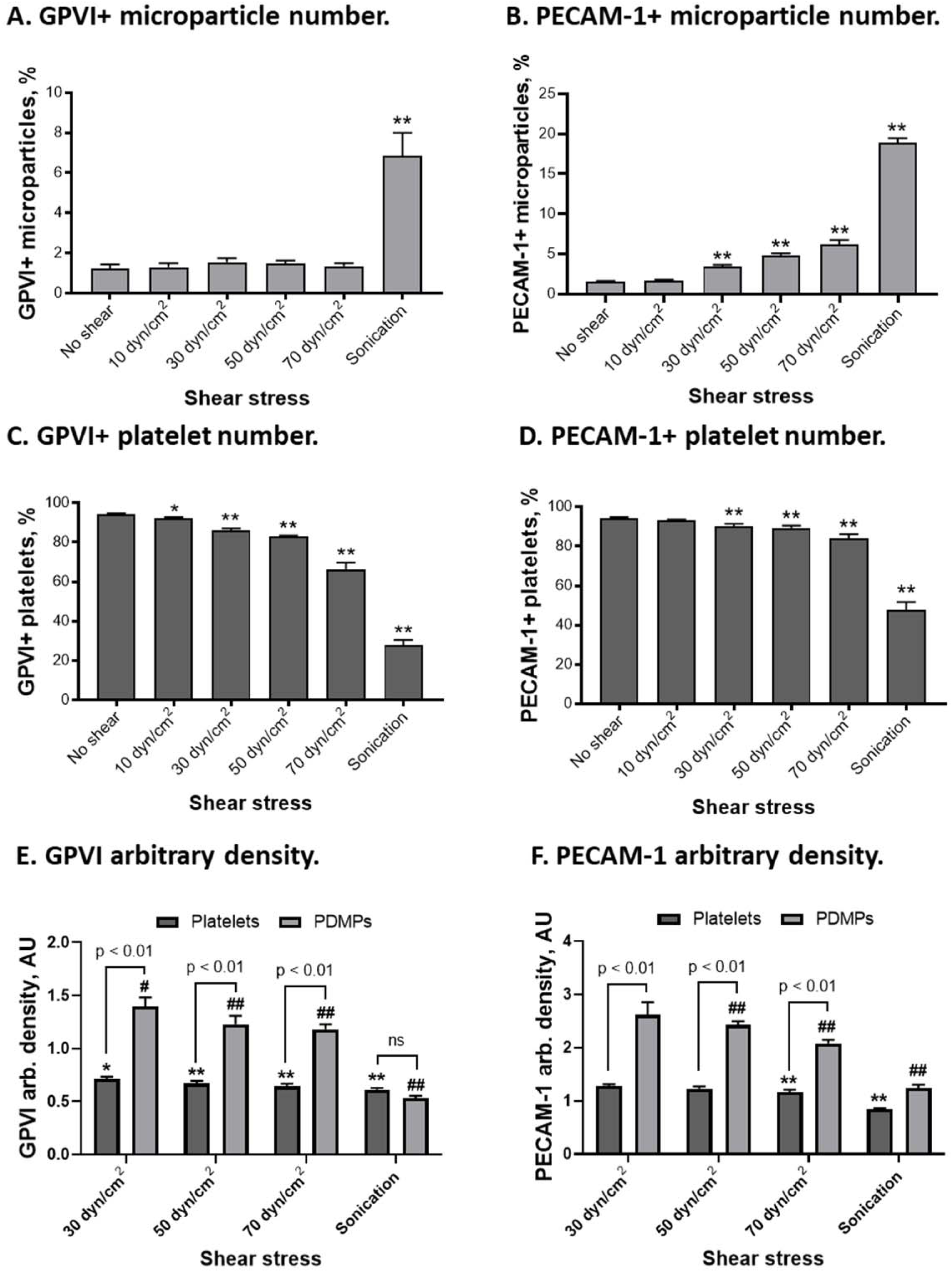
Distribution of GPVI and PECAM-1 receptors on platelets and platelet-derived microparticles (PDMPs): A, B – GPVI+ and PECAM-1+ PDMP number, C,D – GPVI+ and PECAM-1+ platelet number, E, F – arbitrary density of GPVI and PECAM-1 on platelets and PDMPs. N = 6-7. Mean ± SEM, 1-way ANOVA followed by Dunnett multiple comparisons test: *, # - *p*<0.05 and **, ## - *p*<0.01 vs no shear for platelets and PDMPs.

Interestingly, the number of platelets expressing GPVI steeply declined with the increased magnitude of shear exposure, reaching as low as 66.4±3.4% after 70 dyne/cm^2^ shear (**Fig. 5C**). The number of CD31+ platelets also gradually decreased with increasing shear stress yet reaching only 84.0±2.2% after 70 dyne/cm^2^ shear (**Fig. 5D**). Sonication resulted in a significant drop of both platelet populations down to 47.8±3.9% for CD31+ platelets and 28±2.5% for GPVI+ platelets.

Analyzing alteration of platelet size and receptors density on GPVI+ platelets following shear exposure, we did not detect a decrease in platelet size, yet their fluorescence intensity steeply declined with increasing shear stress (**Fig. S4A-B**). As such, the GPVI arbitrary density on platelets significantly decreased with the increased levels of applied shear (**Fig. S4E**). A significant decrease of CD31+ platelet size was noted only following 70 dyne/cm^2^ shear, which offset a slight decrease of CD31 fluorescence intensity, thus indicating a decrease of the receptor arbitrary density on platelets (**Fig. S5A, B, & E**). In contrast, sonication resulted in a significant decrease in platelet size and a 20-25% decrease in GPVI and PECAM-1 arbitrary density on platelets (**Fig. S4E & S5E**, “Sonication”). We also noticed that shear-mediated microparticles were similar in size across all shear conditions and were slightly larger than those generated by sonication (**Fig. S4D & S5D**), while expressing significantly higher levels of GPVI and PECAM-1 on their surface (**Fig. S4F & S5F)**.

When comparing GPVI and PECAM-1 arbitrary density on platelets and microparticles, we found that while not numerous, sheared PDMPs expressed slightly higher levels of GPVI and CD31 on their surface than sheared platelets (**Fig. 5E & 5F**). As such, after 70 dyne/cm^2^ shear, the GPVI and PECAM1 arbitrary density on PDMPs was 1.8× higher than on platelets. Yet again, microparticles generated by sonication demonstrated very similar receptor levels to those on sonicated platelets: 0.5±0.0 AU vs. 0.6±0.0 for GPVI, and 1.2±0.1 AU vs 0.8±0.0 AU for PECAM1, respectively.

### 1.4. P-selectin and PSGL1 distribution on platelets and platelet-derived microparticles

The number of P-selectin+ and PSGL1+ PDMPs increased slightly following platelet exposure to shear stress (**Fig. 6A-B**). In contrast, sonication resulted in a significant increase of both PDMP populations up to 7.2±0.5% for P-selectin+ and 6.2±0.3% for PSGL1+ PDMPs. The baseline levels of both P-selectin and PSGL-1 on platelets were very low, as indicated by the low number of CD62P+ and PSGL1+ platelet populations (**Fig. 6C-D**: “No shear”). Only exposure to 70 dyne/cm^2^ shear resulted in a minor increase of P-selectin+ platelets, yet the number of PSGL1+ platelets remained low across all shear magnitudes tested (**Fig. 6C-D**: “Shear stress”). Sonication resulted in a significant decrease in P-selectin+ and PSGL1+ platelet populations.

**Figure 6.**
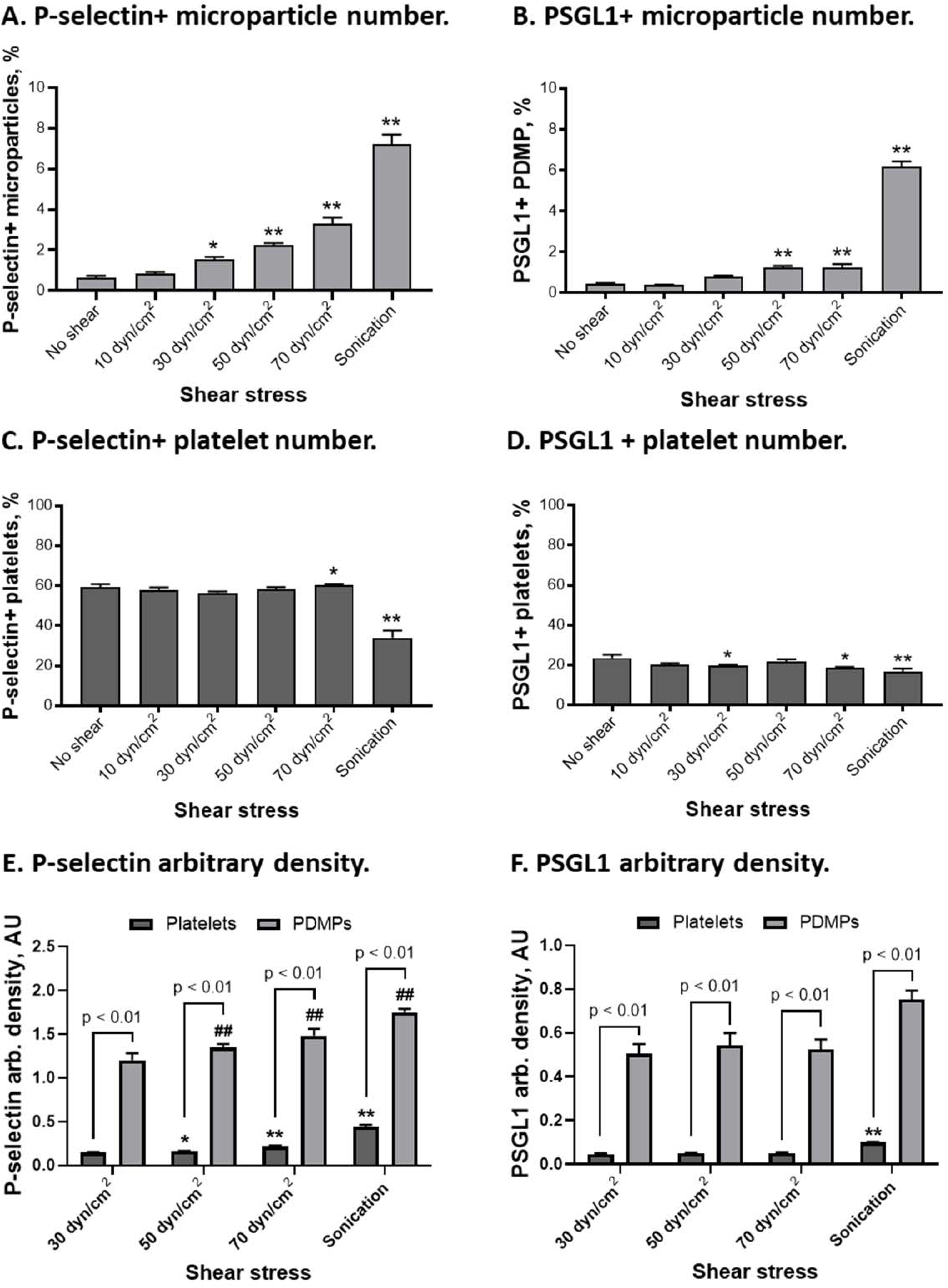
Distribution of P-selectin and PSGL1 on platelets and platelet-derived microparticles (PDMPs): A, B – number of P-selectin+ platelets and PDMPs (CD62+/CD42+ particles), C,D – number of PSGL1+ platelets and PDMPs (CD31+/PSGL1+ particles), E, F – arbitrary density of P-selectin and PSGL1 on platelets and PDMPs. N = 7. Mean ± SEM, 1-way ANOVA followed by Dunnett multiple comparisons test: *, # - *p*<0.05 and **, ## - *p*<0.01 vs no shear for platelets and PDMPs.

Analyzing alterations of platelet size and receptor arbitrary density, we noticed that both high shear and sonication resulted in significant shrinkage of P-selectin+ platelets, yet sonication rendered slightly smaller platelets than shear stress (**Fig. S6B**). The arbitrary density of P-selectin on platelets and PDMPs gradually increased with shear stress magnitude, indicating ongoing degranulation. Sonication resulted in a further increase of P-selectin levels on platelets and PDMPs (**Fig. S6E-F**). Baseline PSGL1 surface expression on platelets was extremely low and remained unchanged across all shear conditions. Interestingly, sonication resulted in a 2-fold increase of PSGL1 density on platelets, but not on PDMPs (**Fig. S7E-F**).

While comparing the receptor arbitrary density on platelets and microparticles, we found that PDMPs generated by shear and sonication yet again expressed significantly higher P-selectin and PSGL1 receptor levels than platelet counterparts (**Fig. 6E-F**). As such, the arbitrary density of P-selectin and PSGL1 on shear-generated PDMPs was 7-10× higher than on sheared platelets. While sonicated PDMPs were less densely populated, but still with a 4-7-fold higher receptor density than platelets.

### 1.5. P2Y12 & PAR1 distribution on platelets and platelet-derived microparticles

PDMPs generated by shear stress and sonication expressed an increased amount of both P2Y12 and PAR1 receptors on their surface (**Fig. 7A-B**). As such, the number of P2Y12+ and PAR1+ PDMP populations gradually increased with the shear magnitude reaching 3.5±0.6% for P2Y12+ and 4.3±0.5% for PAR1+ PDMPs after 70 dyne/cm^2^ shear. Sonication resulted in an even further increase of receptor-positive microparticle populations: 9.6±1.7% and 14.8±1.6% for P2Y12+ and PAR1+ PDMPs, respectively. Analyzing baseline levels of P2Y12 and PAR1 on platelets, we noticed that not all platelets expressed detectable levels of these markers, and their baseline levels were extremely low. As such, only 74.7±1.7 of CD41+ platelets were positive for P2Y12, and 74.0±1.1% of CD41+ platelets were positive for PAR1. Exposure to shear resulted in a minor decrease of both P2Y12+ and PAR1+ platelet populations even following 30 dyne/cm^2^ shear, reaching 63.3±1.7% and 60.7±1.6%, respectively, following 70 dyne/cm^2^ shear. Herein, sonication led to a 3-fold decrease of receptor+ platelet populations (**Fig. 7C-D**, “Sonication”).

**Figure 7.**
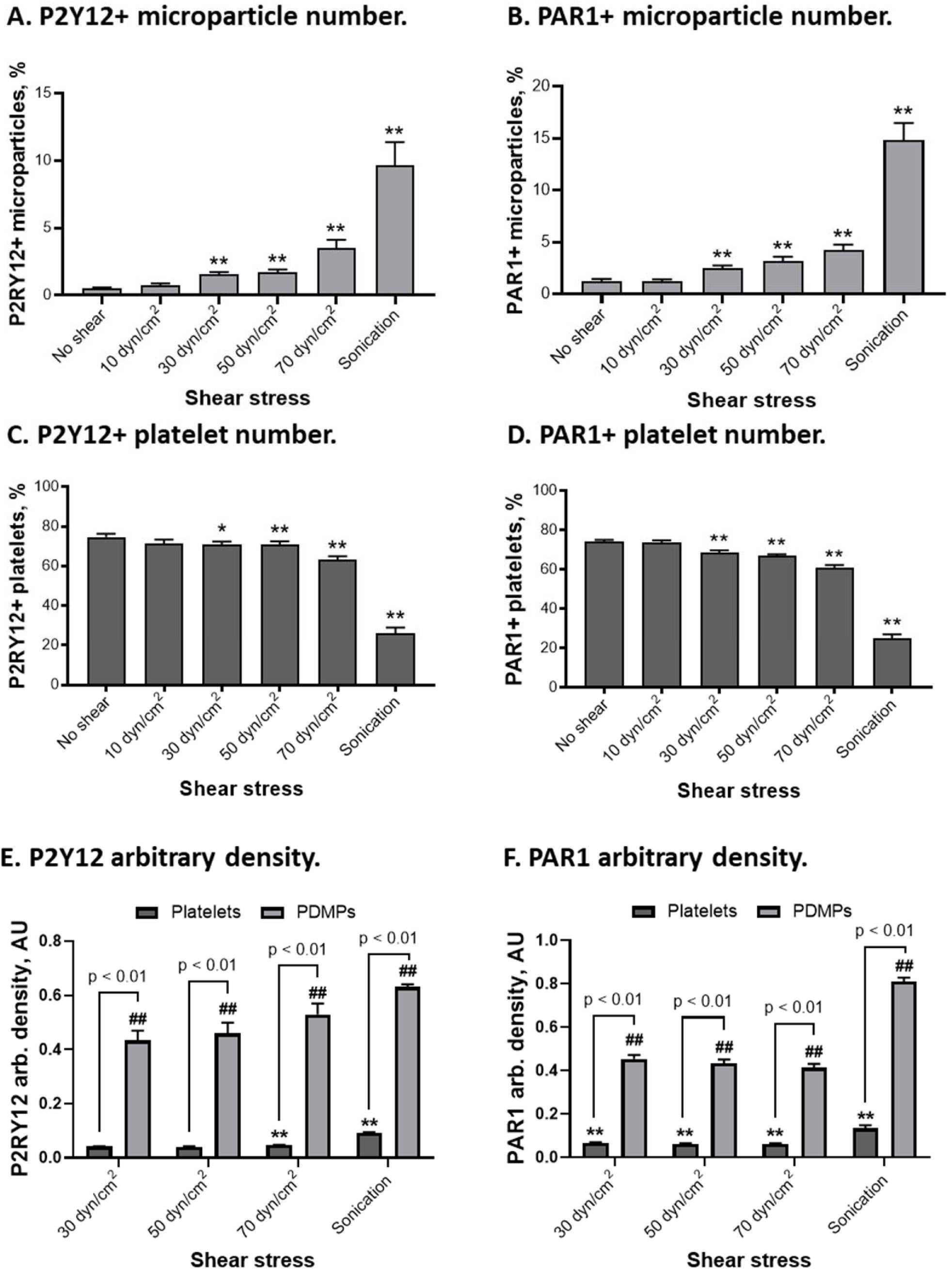
Distribution of P2Y12 and PAR1 on platelets and platelet-derived microparticles (PDMPs): A, B – the number of P2Y12+ platelets and PDMPs (P2RY12+/ CD41+ particles), C,D – number of PAR1+ platelets and PDMPs (PAR1+/CD41+ particles), E, F – arbitrary density of P2Y12 and PAR1 on platelets and PDMPs. N = 4-7. Mean ± SEM, 1-way ANOVA followed by Dunnett multiple comparisons test: **, ## - *p*<0.01 vs no shear for platelets and PDMPs.

Shear stress did not significantly affect platelet size, while sonication rendered significantly smaller platelets than those in unsheared and sheared groups (**Fig. S8B & S9B**). The PDMP size varied across shear levels and sonication (**Fig. S8D & S9D**). Baseline P2Y12 fluorescence intensity and arbitrary density were extremely low and increased slightly following 70 dyne/cm^2^ shear exposure (**Fig. S8A & S8E**), while PAR1 fluorescence and arbitrary density on platelets tended to decrease with the increasing shear stress magnitude (**Fig. S9A & S9E**). Sonication resulted in a nearly 2-fold increase of receptors’ arbitrary density on platelets, but not on PDMPs (**Fig. S8E & S9E**, “Sonication”).

Comparing the receptor density on sheared platelets and PDMPs, we found that PDMPs expressed nearly 10-fold higher levels of P2Y12 receptor and 7-fold higher levels of PAR1 (**Fig. 7E-F**). The PDMP populations generated by sonication were even more enriched with the agonist receptors, expressing 6.5-fold higher levels of P2Y12 and 6.3-fold higher levels of PAR1 than those on sonicated platelets.

### 1.6 Procoagulant activity of platelets and platelet-derived microparticles

The number of platelets and PDMPs binding annexin V significantly increased with the magnitude of applied shear stress (**Fig. 8A-B**). After sonication, the number of annexin V+ platelets was significantly lower than after 70 dyne/cm^2^ shear: 14.3±1.7% vs 21.4±1.9%, respectively. However, sonication rendered a 3-fold higher number of annexin V+ PDMPs: 19.1±1.5% vs 6.0±1.0% for sheared platelets (**Fig. 8B**). Analyzing the density of annexin V binding on platelets exposed to shear and sonication, we found that sheared platelets bound significantly higher levels of annexin V than sonicated counterparts: 4.1±0.0 AU vs 1.4±0.1 AU for 70 dyne/cm^2^ shear and sonication, respectively. Similarly, PDMPs generated by shear stress exhibited 6.2-fold higher levels of annexin V binding than PDMPs generated by sonication (**Fig. 8C**, one-way ANOVA: *p*<0.01). Interestingly, the arbitrary density of annexin V binding on sheared PDMPs was 3-fold higher than on sheared platelets (**Fig. 8C.** PDMPs: 70 dyn/cm^2^ vs.

**Figure 8.**
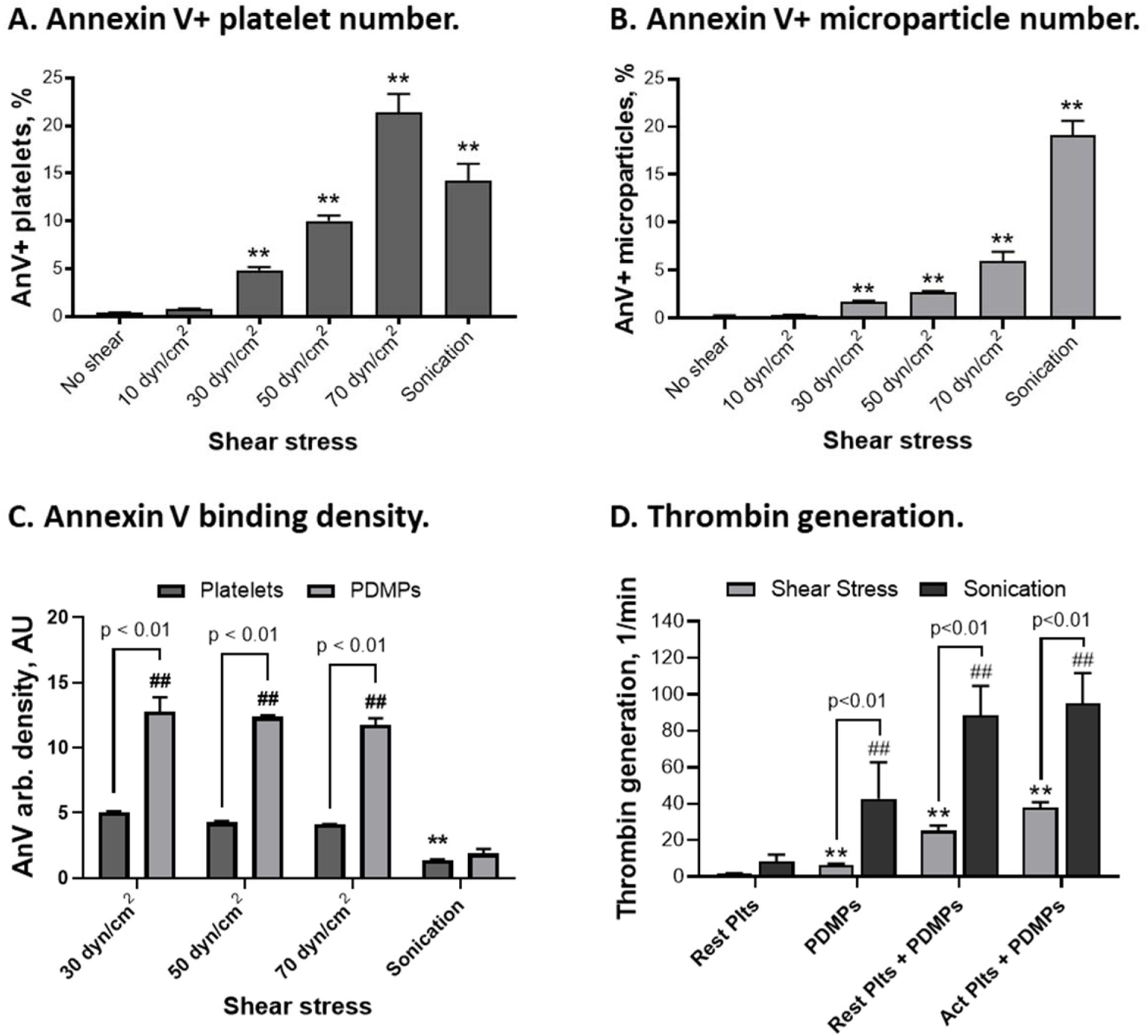
Platelets and platelet-derived microparticles (PDMPs) expose anionic phospholipids on their surface and promote thrombin generation exposure to shear stress and sonication: A, B – the number of platelet and PDMP binding annexin V (AnV+/CD41+ particles), C – arbitrary density of annexin V binding on platelets and PDMPs. D – thrombin generation on platelets and microparticles generated via shear stress exposure (70 dynes/cm^2^, 30 min) or sonication: “Act Plts + PDMPs” – platelets and microparticles following shear exposure or sonication. N = 5-7. Mean ± SEM, 1-way ANOVA followed by Dunnett multiple comparisons test: **, ## - *p*<0.01 vs no shear for platelets and PDMPs.

Platelets: 70 dyn/cm^2^, one-way ANOVA: *p*<0.01), while PDMPs generated by sonication bound the same levels of annexin V as sonicated platelets (**Fig. 8C.** PDMPs: Sonication vs. Platelets: Sonication, one-way ANOVA: *p*>0.05).

To verify the effect of PDMPs on activation of prothrombin by factor Xa, PDMP-rich fractions were obtained from platelets exposed to 70 dyne/cm^2^ shear stress or sonication. We found that platelet exposure to shear stress and sonication promoted thrombin generation, with a resulting thrombin generation rate of 37.9 ± 2.9 min^-1^ and 94.9 ± 16.7 min^-1^, respectively. The PDMPs generated by shear stress showed mild prothrombotic activity when incubated alone or with resting platelets (**Fig. 8D**, one-way ANOVA: *p* < 0.01 vs. “Rest Plts”). Yet, PDMPs generated by sonication rendered a sharp increase in thrombin generation when added alone or with resting platelets, with the resulting thrombin generation rate of 42.7 ± 19.9 min^-1^ and 88.4 ± 2.9 min^-1^, respectively, as compared to 8.2 ± 3.8 min^-1^ in the control group.

### 1.7 The effect of platelet-derived microparticles on platelet aggregation

Under conditions tested, all biochemical agonists induced platelet aggregation, though the aggregation amplitude and area under the curve varied significantly across agonists. In our experimental conditions, ADP and collagen were nearly 10-times as effective in stimulating aggregation as TRAP-6 (**Fig. 9**). We also found that the PDMP-rich fractions added to PRP did not promote platelet aggregation in the absence of aggregation agonist (**Fig. S10**). The extent of the modulatory effect of the two PDMP-rich fractions differed for all agonists tested. Platelet aggregation induced by collagen was significantly inhibited by sheared PDMPs, while sonicated PDMPs showed a less prominent inhibitory effect (**Fig. 9A-B**). As such, in presence of sheared and sonicated PDMPs, collagen-induced platelet aggregation reached only 64.3% and 78.6% of its control level. ADP-induced aggregation was more resilient to inhibition with PDMPs, however shear-mediated PDMPs again demonstrated more potent inhibition than those generated by sonication (**Fig. 9A,C**). Lastly, PDMP-rich fractions did not significantly alter TRAP-6-mediated platelet aggregation. Though, both sheared and sonicated PDMP fractions tended to decrease aggregation as compared with the control group (**Fig. 9A,D**).

**Figure 9.**
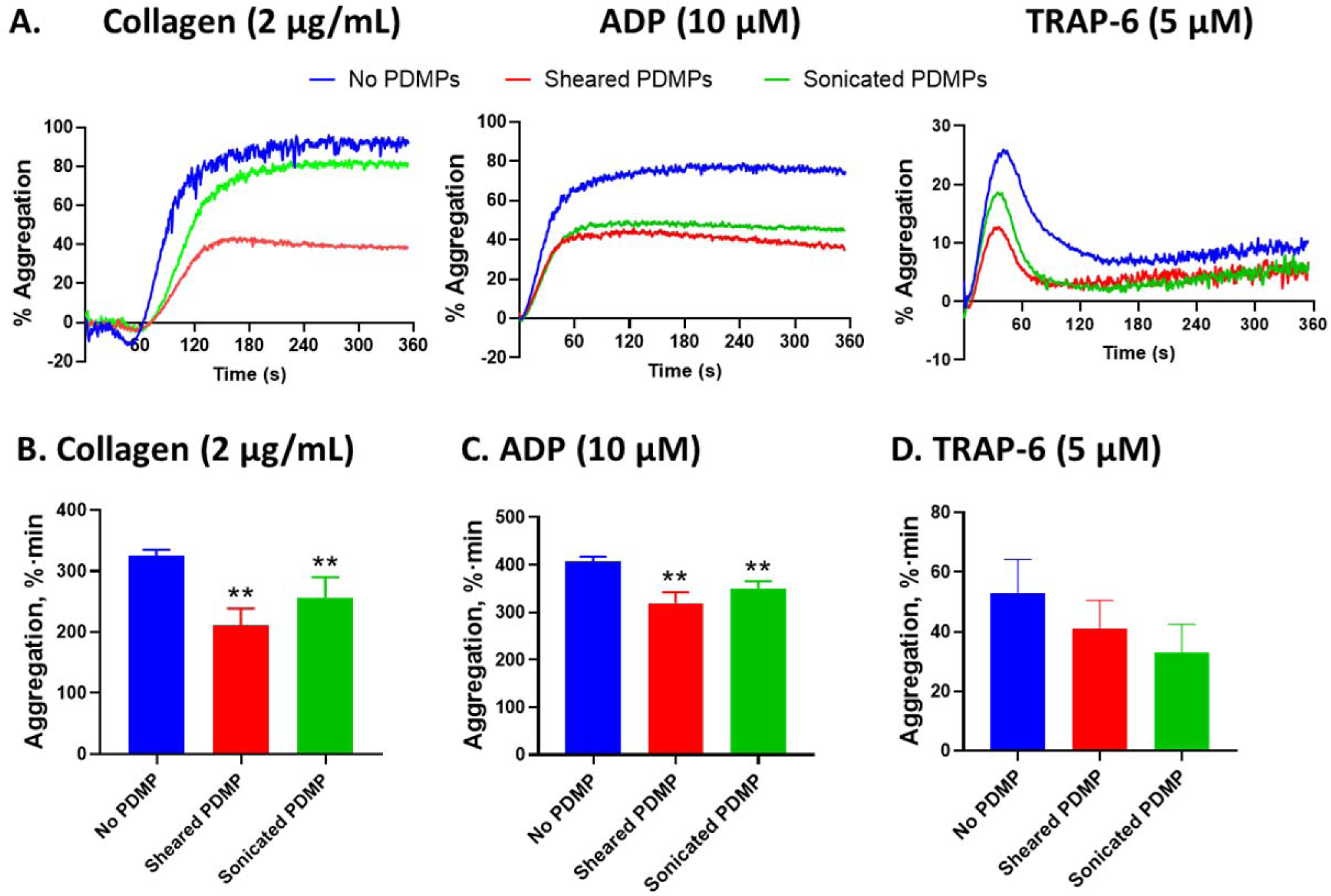
Platelet-derived microparticles (PDMPs) generated as a result of shear stress exposure and sonication inhibit agonist-induced platelet aggregation in plasma: A – representative aggregation curves; B – collagen-induced aggregation (N = 7-14), C – ADP-induced aggregation (N = 7-18), D – TRAP6-induced aggregation (N = 4-10). Mean ± SEM, 1-way ANOVA followed by Dunnett multiple comparisons test: **- *p*<0.01 vs no PDMP.

## DISCUSSION

Persistent exposure to mechanical stresses imparted by implantable CTDs promotes platelet dysfunction and device-related coagulopathy manifested clinically as bleeding and thrombotic adverse events. Our group and others have demonstrated that shear-mediated platelet dysfunction is associated with pro-apoptotic platelet behavior,^36, 77, 100^ decreased platelet aggregation response to biochemical agonists,^37, 101^ and extensive microvesiculation.^37, 76, 77^ Persistently elevated levels of circulated microparticles are associated with adverse events and reduced long-term efficacy of CTD therapy. Here, we defined and characterized the phenotype and functional significance of shear-ejected PDMPs. Using TEM, we demonstrated that exposure to continuous high shear stress or sonication resulted in notable alterations in platelet morphology including shape change, filopodia extension, intense degranulation, and release of PDMPs with varying morphologies, densities, and internal characteristics. Using immunostaining and multi-colored flow cytometry, we found that shear-mediated microvesiculation was associated with differential remodeling of the surface receptors expressed on platelets and PDMPs, with several distinctive patterns emerging. (1) For α_IIb_β_3_ and GPIX adhesion receptors, the arbitrary density on sheared platelets was largely unaltered; yet all sheared PDMPs showed a 2-6-fold higher arbitrary density of these receptors as compared with sheared platelets. (2) For PECAM-1 and GPVI, surface density on platelets and PDMPs gradually decreased with increasing shear magnitude; only a small population of shear-mediated PDMPs was PECAM-1+, but no GPVI+ PDMPs were released with shear exposure. (3) For P-selectin, but not PSGL1, arbitrary density increased slightly on sheared platelets and PDMPs; a small population of shear-mediated PDMPs was P-selectin+ and PSGL1+, expressing significantly higher levels of these receptors in comparison to sheared platelets. (4) The arbitrary density of agonist-evoked receptors P_2_Y_12_ and PAR1 slightly increased on sheared platelets, and only a minor population of shear-ejected PDMPs was positive for P_2_Y_12_ and PAR1, expressing much higher levels of receptors than sheared platelets. (5) PDMPs generated by shear stress had a bidirectional effect on platelet hemostatic function. On the one hand, sheared PDMPs exhibited a negatively charged procoagulant surface promoting thrombin generation; on the other hand, the same PDMPs had an anti-aggregatory effect, inhibiting platelet aggregation in plasma.

Microvesiculation with resultant PDMP formation occurs when platelets undergo hyper-activation associated with a persistent increase in intracellular calcium or enter terminal stages of apoptosis. *In vitro*, hyper-activation can be achieved when platelets are exposed to a combination of strong physiological agonists, e.g. collagen and thrombin, or synthetic agonists as calcium ionophores that instantly increase intracellular calcium, promote cytoskeleton rearrangements, membrane scrambling, and release of procoagulant PDMPs. We and others have previously shown that platelet exposure to device-related shear stress also results in generation of PDMPs.^37, 42, 76, 77^ As an extension of that work, here we showed that platelet exposure to continuous shear stress in free flow is accompanied by notable alterations in platelet morphology including shape change, filopodia extension, quantitative and qualitative alterations of their organelles, and release of extracellular vesicles, ranging from microparticles to exosomes. Following exposure to moderate continuous shear stress (50 dyne/cm^2^), platelets assumed an asymmetrical shape with multiple filopodia extensions, which drastically differed from the round or discoid shape of resting platelets (**Fig. 1 & 2**). In sheared platelets, the number of secretory granules greatly decreased, with the remaining granules grouped together and closely colocalized with the open canalicular system, indicating ongoing degranulation.^98, 102^ It was previously reported that similar morphological changes accompanied PDMP formation in response to activation by strong biochemical agonists.^103^ Microvesiculation induced by thrombin and calcium ionophore was associated with filopodia formation, dilatation of the open canalicular system, degranulation, and cell defragmentation.^103–105^ Our findings are also in agreement with previous studies of increased levels of shear. Shear rate of 10,000 s^-1^ (∼100 dyne/cm^2^) was shown to significantly increase PDMP production in the presence of GPIbα-vWF interaction, with drop-like or spherical shaped particles formed from detached membrane tethers extending downstream of adhered platelets.^106^ Similarly, increased generation of PDMPs was demonstrated when whole blood was exposed to prolonged shear stress *ex vivo* in an ECMO circuit.^107^ The number PDMPs increased with the circulation time, suggesting a role of stress accumulation in promoting persistent PDMP formation.^107^

Here, we observed that exposure to supraphysiologic shear resulted in significant alterations of platelet morphology, with only two-thirds of platelets remaining whole and intact. Many platelets resembled a ghost-like appearance with pale dispersed cytoplasm occupied by numerous membrane-enclosed vesicles (**Fig. 2B-C**). This vesicles are likely pro-PDMPs or multivesicular PDMPs similar to those reported for thrombin-induced activation.^103^ Platelet exposure to sonication resulted in near complete platelet disruption with only a few remaining ghost-like platelet structures. Sheared PDMPs were numerous and substantially varied by size, density, and morphology. In contrast, PDMPs generated by sonication were largely uniform in size though demonstrated significant heterogeneity of density and structure, ranging from dark dense particles to seemingly transparent hollow-shell structures.

The morphological heterogeneity of PDMPs generated by shear and sonication suggests that a variety of mechanisms may be operative in PDMP formation. Precedent for a range of operative mechanisms and morphologic heterogeneity of PDMPs exists with strong biochemical agonists. It has been reported that thrombin and calcium ionophore promote formation of four ultrastructurally different types of microvesicles: single membrane-enclosed vesicles, multilayer vesicles, multivesicular particles consisting of several vesicles (10-15), and particles with dense contents including cytoplasm and organelles of parental cells.^103^ It has been suggested that these four types of PDMPs are formed as a result of differing mechanisms including membrane invagination following budding; formation at the end of pseudopodia; via degranulation involving the open canalicular system; or platelet fragmentation. While these mechanisms may be operative in shear-mediated PDMP generation, the morphologic observations in our study are also consistent with a previously proposed mechanodestruction mechanism wherein shear stress imparts direct mechanical damage to the platelets resulting in enhanced membrane porogenicity, rapid influx of calcium and other ions, and ultimate membrane fragmentation generating microparticles and exosomes.^35^ The mechanodestruction mechanism is supported by prior modeling studies of membrane and whole cell shear stress accumulation,^108–111^ and is consistent with our present TEM observations. In summary, our ultrastructural study of platelet microvesiculation following shear exposure allows us to conclude that (1) sheared platelets undergo shape change with cytoplasmic extrusion and pseudopodia extension; (2) along with PDMP generation, sheared platelets are engaged in extensive degranulation as evidenced by a decrease in the number of secretory granules and dilatation of the open canalicular system; (3) not all platelets are actively involved in PDMP generation, with some platelets exhibiting the morphology of intact cells, while others sustain damage resulting in ghost-like membrane structures packed with pro-PDMPs; (4) shear-mediated PDMPs vary in size, density, and ultrastructural features, suggesting that a range of mechanisms of microparticle formation are involved. The multiplicity of mechanisms operative in shear-mediated microparticle formation may include specific biochemical means involving mechanotransduction pathways and direct cell and membrane damage via mechanodestruction.

Platelet surface receptors are responsible for sensing biochemical and mechanical signals from the intravascular environment, transducing these signals inside the cell, and initiation a functional response. As the name suggests, *adhesion receptors* are crucial for platelet adhesion to vascular walls and aggregation to one another.^50, 51^ As platelets mature and age, the basal levels of receptors’ surface expression change. Platelet maturation is associated with an increase of basal surface expression, while platelet aging and apoptosis result in downregulation of receptor surface expression. Enzymatic shedding and microvesiculation have been identified as two primary mechanisms responsible for the downregulation of platelet surface receptors. Our research group was one of the first to recognize that platelet exposure to continuous shear stress of increased magnitude and duration promotes downregulation of GPIb, α_IIb_β_3_, and P-selectin surface expression via generation of PDMPs carrying increased levels of these receptors.^37, 112^ Herein, we conducted a quantitative analysis of shear-generated PDMPs under differing shear conditions and characterized the surface expression of six types of adhesion receptors (α_IIb_β_3_, GPIX, GPVI, PECAM-1, P-selectin, PSGL-1) on sheared platelets and PDMPs.

Platelet α_IIb_β_3_ and GPIb-IX receptor complexes are abundantly expressed on the platelet surface and have been extensively studied as targets for antiplatelet therapeutics as the primary molecular machinery platelets rely on during adhesion and aggregation. Binding to its primary ligand vWF, GPIb-IX was also identified as a platelet shear sensor and anchor reinforcing platelet aggregation under shear.^113^ Analyzing PDMPs carrying integrin α_IIb_β_3_ and GPIX on their surfaces, we found that continuous exposure to shear stress up to 70 dyne/cm^2^ induced a 5-fold increase in the number of both populations of circulating PDMPs, though the GPIX+ PDMPs were less numerous than their α_IIb_β_3_+ counterparts. A slight decrease of CD41+ and CD42a+ platelets was also registered, likely associated with platelet disintegration accompanying PDMP generation, as reflected in our TEM observations. Shear-mediated PDMPs were increasingly decorated with both α_IIb_β_3_ and GPIX (**Fig. 4E-F**). These findings are in agreement with our previous report, where we first demonstrated the shear-mediated generation of PDMPs carrying increased levels of platelets receptors α_IIb_β_3_ and GPIb following exposure to continuous but not pulsatile shear of increased magnitudes.^37^ Interestingly, the surface density of GPIX, but not α_IIb_β_3_, on sheared PDMPs was slightly elevated with increasing shear magnitude. The upregulation of GPIX expression on PDMPs may result from redistribution of internal GPIX from platelet granules following their fusion with platelet plasma membrane or open canalicular system when granule exocytosis occurs prior to or coincides with microvesiculation. Redistribution of GPIb and GPIX between platelet surface and internal membranes was previously demonstrated during platelet stimulation with strong biochemical agonists and neutrophil proteinases.^114^ Alternatively, the GPIX-enriched PDMP population may entirely originate from platelet internal granules carrying significant amounts of GPIb-IX,^115, 116^ as degranulation intensifies with increasing shear magnitude. Platelet sonication, used as a positive control for platelet destruction caused by mechanical forces, resulted in platelet disintegration as indicated by a significant platelet count drop and generation of increased amounts of PDMP-sized bodies, carrying both α_IIb_β_3_ and GPIX receptors. However, the surface density of α_IIb_β_3_ and GPIX on microvesicles generated by sonication remained the same or comparable to the receptor density on platelets (**Fig. 4E-F**). The last observation strongly suggests that sonication nonspecifically destroys platelets, while shear stress induces a specific platelet response associated with selective redistribution of platelet receptors from platelets to PDMPs and is likely accompanied by platelet degranulation.

Platelet GPVI and PECAM-1 receptors, while not as abundant as α_IIb_β_3_ and GPIb-IX-V, play a prominent role in platelet rolling and adhesion to vascular endothelium, regulation of primary hemostasis and thrombus formation. GPVI is also regarded as a potent signaling receptor for collagen-evoked platelet activation and microparticle release when co-stimulated along with other potent biochemical agonists, i.e. thrombin and ADP.^117, 118^ An increasing body of evidence suggests that PECAM-1 is involved in the negative regulation of thrombus formation.^119^ Stimulation of PECAM-1 has been shown to inhibit platelet activation mediated by a range of receptors, including GPVI, GPIb-V-IX, and G-protein coupled receptors, while PECAM-1 knockout resulted in increased platelet sensitivity to agonists and hyper-aggregability.^119–121^ We found that following shear exposure, surface expression of GPVI and PECAM-1 on platelets significantly decreased as indicated by the incremental decrease of their arbitrary density. The number of GPVI+ and PECAM-1+ platelets also decreased, which might be attributed to the downregulation of receptors’ surface expression and platelet disintegration as a result of microvesiculation. We found that sheared PDMPs did not express GPVI, while a small population presented PECAM-1 on their surface (**Fig. 5A-B**). The PECAM-1 density on PDMPs was higher than on sheared platelets and tended to decrease with the magnitude of shear applied. Proteolytic shedding of both GPVI and PECAM-1 associated with activation, apoptosis, and exposure to hypershear stress has been extensively documented for both platelets and endothelial cells.^62, 122, 123^ It was previously shown that microvesicles released following platelet activation by biochemical agonists also contain PECAM-1, along with integrins α_IIb_β_3_ and β_1_. Our observations suggest that following shear exposure both GPVI and PECAM-1 are downregulated on platelets, while PECAM-1, but not GPVI, is found on shear-mediated microparticles where it is also downregulated (**Fig. 5F**). In contrast, sonication resulted in a massive generation of PECAM-1+ and GPVI+ PDMPs and a corresponding decrease of platelet populations carrying those receptors. Following sonication, the surface density of those receptors was virtually the same on PDMPs and platelets (**Fig.5E-F**), strongly suggesting that sonication induced random platelet fragmentation rather than a controlled process of PDMP formation.

Of the six adhesion receptors characterized in this study, P-selectin and PSGL-1 are the only receptors whose surface expression on platelets is inducible. In resting platelets, P-selectin is located within α-granule membranes. Upon activation and degranulation, P-selectin is translocated onto the platelet surface.^124^ Via P-selectin, platelets interact with leukocytes and endothelial cells promoting their activation, expression of pro-inflammatory proteins, and microvesiculation. When underutilized, P-selectin undergoes rapid “shutdown” by shedding of a soluble fragment (sP-selectin) from the platelet surface or internalization.^124, 125^ Similarly, the reported level of PSGL-1 expression on unstimulated platelets is very low (25-100× less than on leukocytes), increasing following platelet activation with thrombin, and gradually decreasing as platelets age.^50^ Examining the distribution of P-selectin and PSGL-1 on platelets under shear, we also found that unstimulated platelets expressed a very low level of both receptors. The surface density of P-selectin, but not PSGL-1, mildly increased with exposure to increasing shear magnitude, indicating ongoing degranulation. A small population of shear-ejected PDMPs expressed increased levels of both P-selectin and PSGL-1 (**Fig. 6E-F**). A detected mild increase of P-selectin on sheared platelets might be indicative of low levels of secretory activity stimulated by shear. Yet, we believe that seemingly low levels of P-selectin on sheared platelets are rather a snapshot of transient P-selectin exposure on platelets, followed by rapid shedding of its soluble fragment from the platelet surface. We have previously demonstrated that platelet exposure to MCS-generated shear stress for an extended period of time is associated with increased levels of circulating sP-selectin *in vitro* and *ex vivo*.^37, 112^ Increased levels of P-selectin on PDMPs also supports this hypothesis, suggesting that platelet degranulation occurs prior to or simultaneously with PDMP generation. P-selectin is preserved on PDMP surface, possibly due to the lower extent of shedding on PDMPs as compared to platelets. Increased levels of P-selectin expression were previously reported for PDMPs released by platelets activated with thrombin.^126^ Whether a similar mechanism of transient expression and subsequent shedding from the platelet surface is operative for PSGL-1 remains unclear. Yet, it is noteworthy that both P-selectin and PSGL-1 were abundantly expressed on microvesicles generated by platelet sonication (**Fig. 6A-B**). This observation further confirms the random nature of platelet destruction and loss of integrity caused by sonication, with the resultant release of platelet secretory granules carrying high levels of both P-selectin and likely PSGL-1. Indeed, our TEM imaging of sonicated PDMPs visually identifies several populations of optically dense vesicles resembling the appearance of secretory granules previously seen inside unstimulated platelets (**Fig. 1H**).

The most well-known and therapeutically relevant *agonist receptors* are ADP-evoked P_2_Y_12_ and thrombin-evoked PAR1 receptors. The surface expression of these receptors is differentially regulated over the platelet lifespan correlating with platelet activation response.^127^ Upregulation of P_2_Y_12_ receptor on the platelet surface and augmented mRNA expression was recently described in patients with acute coronary syndrome, type 2 diabetes mellitus, and sepsis, as associated with platelet hyperreactivity and thrombosis.^128–130^ The surface expression of PAR1 receptors widely varies among healthy individuals^131^ and was shown to increase in various inflammatory conditions, including atherosclerotic cardiovascular disease, arthritis, and asthma.^132, 133^ Analyzing the surface expression of P_2_Y_12_ and PAR1 on platelets under shear, we found that levels of these receptors on resting platelets are rather low; both P2Y12+ and PAR1+ platelet populations further decreased with increasing shear magnitude (**Fig. 7C-D**). The surface density of P_2_Y_12_ on platelets remained unchanged, while PAR-1 decreased slightly following shear exposure. Similar to adhesion receptors, the downregulation of agonist-evoked receptors on platelets might be a result of plasma membrane reorganization and receptor redistribution into sheared PDMPs. Indeed, small P_2_Y_12_+ and PAR1+ populations of sheared PDMPs were detected in our study (**Fig. 7A-B**), and such a possibility was discussed by others earlier.^127^ Alternatively, it has been shown that platelet hyper-activation by biochemical agonists is associated with desensitization and hence internalization of P_2_Y_12_ and PAR1 receptors.^131^ The internalization of a partially cleaved form of PAR-1 by proteinases other than thrombin was also reported.^132^ The internalized pool of these receptors was further tracked down to platelet endosomes and lysosomes.^134^ We do not exclude the possibility that a small PDMP population carrying both P_2_Y_12_ and PAR1 receptors and showing up to 10-fold higher levels than sheared platelets might originate from platelet endosomes. Indirect confirmation of this hypothesis yet again comes from sonicated platelets and PDMPs. Unlike shear, sonication resulted in a substantial decrease of both P_2_Y_12_+ and PAR1+ platelets and a coincidental increase of PDMP populations carrying these receptors (**Fig. 7A-D**). Nearly one-third of CD41+ sonicated PDMPs were loaded with P2Y12 and/or PAR1 receptors. Recent proteomic analysis of PDMPs released by resting and thrombin-activated platelets confirmed the presence of lysosomal proteins and specifically those regulating PAR1 internalization and recycling.^135^

Lastly, we examined the effect of PDMPs generated by shear or sonication on platelet hemostatic function, namely thrombin generation and aggregation. Earlier, we demonstrated that sheared, but not biochemically activated, platelets expose negatively charged phospholipids on their surface and promote thrombin generation.^36^ Here, we found that platelet exposure to mechanical forces, either shear or sonication, promote externalization of phosphatidylserine on the platelet surface, as indicated by increased binding of annexin V, and facilitate shedding of numerous PDMPs that also bind annexin V. The annexin V binding density on sheared PDMPs was nearly 3-fold higher than on sheared platelets (**Fig. 8C**), indicating that the density of negatively charged phospholipids on PDMPs is much higher than on platelets. In contrast, sonicated platelets and PDMPs demonstrated almost identical binding capacity for annexin V which was 6-fold lower than for sheared PDMPs. Sonication also resulted in much higher number of annexin V+ microparticles than those produced by shear stress. Taken together, these observations suggest that differing mechanisms of phosphatidylserine externalization are taking place in sheared and sonicated platelets. Specifically, increased phosphatidylserine surface density on sheared platelets and PDMPs is indicative of intentional and coordinated membrane lipid reorganization likely resulting from preceding signaling events and activation of platelet scramblases. Conversely, low-density annexin V binding on sonicated platelets and PDMPs may result from random platelet membrane damage, fragmentation, and formation of numerous microvesicles, some of which may be inverted with the internal membrane lipid layer enriched with anionic phospholipids being externally exposed.

Interestingly, sheared and sonicated PDMPs demonstrated differing functional effects on thrombin generation and platelet aggregation. Sonicated PDMPs were more potent promoters of thrombin generation than PDMPs released by shear stress when added alone or alongside resting or activated platelets (**Fig. 8D**). We speculate that more abundant sonicated PDMPs cumulatively offer a larger net procoagulant surface, thus resulting in higher rates of thrombin generation than less abundant sheared PDMPs having a higher phosphatidylserine density on their surface. Moreover, demonstrating substantial prothrombinase activity, PDMPs may be more potent that platelets, even after short bursts of high shear stress, with prothrombinase activity persisting and increasing during subsequent low shear regimes.^34^ When added to intact platelets in the plasma environment, both sheared and sonicated PDMPs inhibited platelet aggregation induced by collagen, ADP, and to a lesser extent by TRAP-6. Sheared PDMPs showed a slightly more pronounced anti-aggregatory effect than sonicated PDMPs (**Fig. 9**). As such, our findings indicate that PDMPs generated by mechanical forces similar to those existing within device-supported circulation assume two antagonistic roles, being both procoagulant and antiaggregatory. Densely covered with negatively charged phospholipids, PDMPs promote activation of prothrombin by factor Xa and generation of thrombin. As small and mobile membrane vesicles, PDMPs might also contribute to an acceleration of thrombosis and dissemination of the prothrombotic milieu from the bloodstream into organ tissues, thus contributing to microthrombosis associated with implantable CTDs.^80, 136, 137^ On the other hand, we demonstrated that shear-generated PDMPs are also enriched with adhesion receptors (α_IIb_β_3_, GPIb-IX, GPVI, and PECAM-1) that are redistributed from platelets as a result of shear exposure. We do not exclude the possibility that PDMPs can compete with platelets for binding sites on the platelet surface and for adhesion protein ligands in plasma when platelets aggregate, thus effectively inhibiting aggregation. This observation is indirectly supported by the fact that sheared PDMPs with a higher surface density of major adhesion receptors also showed a more pronounced anti-aggregatory effect as compared with sonicated PDMPs. As such, shear-mediated generation of PDMPs showing an anti-aggregatory effect and associated decrease of adhesion receptors on the platelet surface jointly contribute to device-related bleeding coagulopathy. Our *in vitro* findings are in alignment with the latest reports studying platelet function in device-implanted patients. It was shown that the severity of acquired platelet defects, i.e. reduced GPIbα and GPVI expression and reduced α_IIb_β_3_ activation, is predictive of non-surgical bleeding events in VAD recipients, having even higher predictive power than vWF degradation parameters.^80^ It was also reported that device-implanted patients with increased bleeding risk demonstrate decreased platelet aggregation response to biochemical agonists (ADP, TRAP-6, and arachidonic acid), underscoring the importance of these acquired defects as drivers of device-related bleeding coagulopathy.^138^

Routine flow cytometry used to capture and characterize shear-mediated PDMPs is known to underestimate the small-size microparticle population (<300 nm) and exosomes (<100 nm). Therefore, the number of shear-ejected microvesicles detected in our study might also be underestimated. Using TEM for the visualization of platelets and microparticles requires extensive sample processing. As such, it is rather difficult to visually distinguish circulating PDMPs from platelet filopodia or platelet fragments released as a result of sample processing due to the apparent morphological similarity of these structures.

In summary, our findings indicate that exposure to supraphysiologic shear stress accumulation or sonication promotes notable alterations in platelet morphology including shape change, filopodia extension, intense degranulation, and generation of PDMPs. Shear-mediated microvesiculation is associated with differential remodeling of the surface receptors expressed on platelets and microparticles, with several distinctive trends emerging. (1) The surface density of α_IIb_β_3_ and GPIX adhesion receptors on sheared platelets remains largely unaltered; yet, sheared PDMPs show a significantly higher surface density of these receptors. (2) Following shear exposure, PECAM-1 and GPVI expression on platelets gradually decreased; yet only a small population of shear-mediated PDMPs was PECAM-1 positive, while no GPVI+ PDMPs were detected following shear. 3) The surface density of P-selectin, but not PSGL1, slightly increased on sheared platelets and PDMPs; a small population of sheared PDMPs was P-selectin+ and PSGL1+, expressing significantly higher levels of these receptors than sheared platelets. 4) The surface density of agonist-evoked receptors P_2_Y_12_ and PAR1 slightly increased on sheared platelets, with only a minor population of sheared PDMPs being positive for P_2_Y_12_ and PAR1, expressing much higher levels of receptors than sheared platelets. Lastly, we found that PDMPs generated by shear stress had a bidirectional effect on platelet hemostatic function. On one hand, sheared PDMPs exhibit a negatively charged procoagulant surface and promote thrombin generation; on the other hand, PDMPs have an anti-aggregatory effect, inhibiting platelet aggregation in plasma induced by collagen and ADP. Which mechanism prevails *in vivo* may vary in individual circumstances and remains to be defined. Though, it has become clear that PDMP generation is a selective and regulated process associated with the alteration of platelet morphology and function, and could be considered as both cause and effect of shear-mediated platelet dysfunction.

From a translational perspective our findings of heterogeneity in platelet response to shear and significant differences in PDMP phenotype, underscores that a range of operative mechanisms are at play in the platelet response to supraphysiologic shear imparted by CTDs. When coupled with the recognition of reduced efficacy of present antiplatelet agents as means of limiting CTD adverse events, this work supports the need to develop novel pharmacologic strategies that effectively target shear-mediated platelet dysfunction and generation of PDMPs. Successfully addressing these issues provides a great opportunity for advancing the efficacy of CTDs while reducing patient risk.

## ACKNOWLEDGEMENTS

Y. Roka-Moiia designed and performed experiments, analyzed and interpreted data, wrote and edited the manuscript; S. Miller-Gutierrez performed experiments; K.R. Ammann and J.E. Italiano performed imaging experiments, analyzed data, edited the manuscript; J. Sheriff, D. Bluestein, J.E. Italiano, and R.C. Flaumenhaft participated in discussions and reviewed the manuscript; M.J. Slepian designed research, analyzed imaging data, and wrote and edited the manuscript.

